# Water depletion can generate contrasting root allocation responses to belowground competition

**DOI:** 10.64898/2026.08.09.743548

**Authors:** Ciro Cabal, Mario Chico Rodríguez

**Affiliations:** Department of Biology, Rey Juan Carlos University (URJC), Móstoles, Spain; Global Change Research Institute, Rey Juan Carlos University (IICG-URJC), Móstoles, Spain

**Author notes:** **Statement of authorship:** C.C. designed research; C.C. and M.C.R. performed research; C.C. developed and analyzed the optimization model; C.C. and M.C.R. analyzed data; C.C. administered the project and acquired funds; and C.C. and M.C.R. wrote the paper. **Data availability statement:** The data and R code supporting the findings of this study (Cabal & Chico-Rodríguez 2026) are openly available at https://doi.org/10.5281/zenodo.21863880.

**Keywords:** Belowground competition, Optimization model, Root allocation, Root tragedy of the commons, Soil resource dynamics, Soil water depletion

## Abstract

Plants competing belowground may produce extra roots, fewer roots, or no detectable change compared to plants growing alone. This inconsistency is often attributed to plants altering their root allocation in response to diverse cues, including neighbor detection and resource depletion by neighbors, but isolating these cues experimentally is challenging. Here, we hypothesize that water depletion alone can generate the range of root allocation strategies reported in the literature. We present this hypothesis as a water-explicit optimization model of root allocation that predicts a non-monotonic response. The model identified a critical depletion rate at which allocation shifted from increasing to decreasing with depletion. We tested this prediction using artificially rooted pots that imposed controlled water depletion while excluding living neighbors and their cues. A continuous artificial depletion gradient revealed the predicted hump-shaped pattern. These results reframe root overproliferation and underproliferation as positions along a single depletion-response curve.

## Introduction

Roots acquire water and most mineral nutrients required for plant growth, making root allocation a key determinant of plant productivity, vegetation carbon storage (Farrior *et al*. 2015; Ma *et al*. 2021), and crop yield (Homulle *et al*. 2022). Yet predicting how plants alter root allocation under belowground competition remains a long-standing unresolved problem in plant ecology. Experimental studies have reported root overproliferation, root underproliferation, and no detectable root allocation response to neighbors (Belter & Cahill 2015; Smyčka & Herben 2017; Gottlieb & Gruntman 2024). It has been proposed that this diversity of outcomes arises because plant species may differ in how they respond to a wide array of environmental cues including neighbor detection, self/non-self root recognition, and resource depletion by neighbors (Schenk 2006; Cahill *et al*. 2010; Cahill & McNickle 2011; Novoplansky 2019).

Early game-theoretical work on crop ideotypes proposed that competition for limiting resources can favor redundant investment in competitive organs, including roots in water-limited crops, even when this reduces collective yield (Zhang *et al*. 1999). This represents a root version of the tragedy of the commons, in which individually advantageous resource-foraging strategies reduce collective performance (Hardin 1968). The root tragedy model later formalized this logic for belowground competition, predicting that plants sharing a common resource pool should overproliferate roots relative to plants with exclusive access to their own space (Gersani et al. 2001). Some experiments, seemingly consistent with the root tragedy of the commons, have found that plants increase root production in the presence of neighbors (e.g., Gersani *et al*. 2001; Maina *et al*. 2002; O’Brien *et al*. 2005; Mercer & Eppley 2014). This prediction has often been interpreted as a proactive response to non-self roots, implying that plants overproduce roots because they detect competitors (Schenk 2006; Cahill & McNickle 2011; Semchenko *et al*. 2014), or because their root tips can distinguish whether nearby roots are self or non-self at the plant level (Callaway 2002; Falik *et al*. 2003; Chen *et al*. 2012).

Other studies have presented results that are inconsistent with this prediction, including species-specific responses, root underproliferation, or no detectable change in absolute root allocation (e.g., Cahill 2003; Armas & Pugnaire 2011; Semchenko *et al*. 2014; Chen *et al*. 2015, 2021; Gottlieb & Gruntman 2024; Cabal et al. 2024). Deviations from the tragedy-of-the-commons prediction have been attributed to the complexity of plant behavioral strategies under multiple environmental cues (Cahill *et al*. 2010; Cahill & McNickle 2011), including cases in which plants do not respond to neighbor detection directly but only to reduced nutrient availability (McNickle & Brown 2014). Yet the original tragedy-of-the-commons logic does not necessarily require such neighbor detection. As argued by Cabal (2022), plants modeled by Gersani *et al*. (2001) may simply increase root allocation because neighbors modify soil resource depletion and thereby change the net gains from further root investment.

In conventional additive competition designs, in which neighbors are added to a focal plant, plants grown alone and plants grown with neighbors typically differ not only in the presence of competitors, but also in the total amount of soil resources available per plant, the volume of soil available for exploration, or the concentration of resources in that soil (Hess & De Kroon 2007; Semchenko *et al*. 2007). Pot experiments comparing plants grown alone and with neighbors generally cannot simultaneously control pot volume, total nutrients per plant, and soil nutrient concentration, so at least one of these factors is necessarily confounded with neighbor addition (Chen *et al*. 2020; McNickle 2020). As a result, observed changes in root allocation may reflect responses to neighbors themselves, responses to altered resource availability, or artifacts of experimental design. Some experiments have attempted to isolate neighbor-induced resource-depletion effects from other biological cues through nutrient manipulation (Messier *et al*. 2009; Nord *et al*. 2011). However, from a resource-dynamics point of view, experimentally controlling resource concentration through nutrient addition is not equivalent to varying resource depletion, and the two may trigger different root proliferation responses (Cabal *et al*. 2021). A mechanistic understanding of root allocation therefore requires experimental approaches that dissociate resource depletion from resource addition and neighbor-derived cues.

Modeling developments have partly addressed this problem by formalizing alternative hypotheses about how plants should allocate roots under belowground competition. Models that incorporate soil volume and nutrient availability have clarified whether apparent root overproliferation reflects strategic competition, resource availability, or experimental design (O’Brien & Brown 2008). Other models have contrasted game-theoretical neighbor pre-emption with ideal-free-distribution strategies in which plants allocate roots according to resource availability rather than direct neighbor responses (McNickle & Brown 2012, 2014). However, as in experiments that control resource concentration, these models generally treat resource availability as a fixed amount or patch quality, rather than as a dynamic state variable governed by input, abiotic loss, and depletion by roots. More recent models have incorporated such soil resource dynamics directly, showing that root allocation patterns can emerge from the balance between resource input, abiotic loss, and biotic uptake (Cabal *et al*. 2020, 2021). However, whether depletion dynamics alone can generate contrasting whole-plant root allocation responses remains unresolved.

Here, we developed a mathematical model predicting optimal root allocation as a function of explicit soil water depletion imposed by neighboring roots. The model provides a non-game-theoretical counterpart to root tragedy logic: rather than modeling strategic interactions between two plants, it asks how a single plant should adjust root biomass when a limiting soil resource is depleted at increasing neighbor-equivalent rates. Based on the model predictions, we hypothesize that root allocation should increase under moderate water depletion but decline once depletion exceeds a critical threshold. Under this framework, root overproliferation, no detectable response, and underproliferation can emerge as different positions along a single depletion-response curve, showing that resource depletion dynamics alone can generate contrasting root allocation responses without invoking direct neighbor detection. We tested this hypothesis experimentally using artificially rooted pots (ARPs; **Fig. 1**), which impose controlled rates of soil water depletion in the absence of living neighbors and therefore eliminate neighbor-derived biological cues.

**Figure 1.**
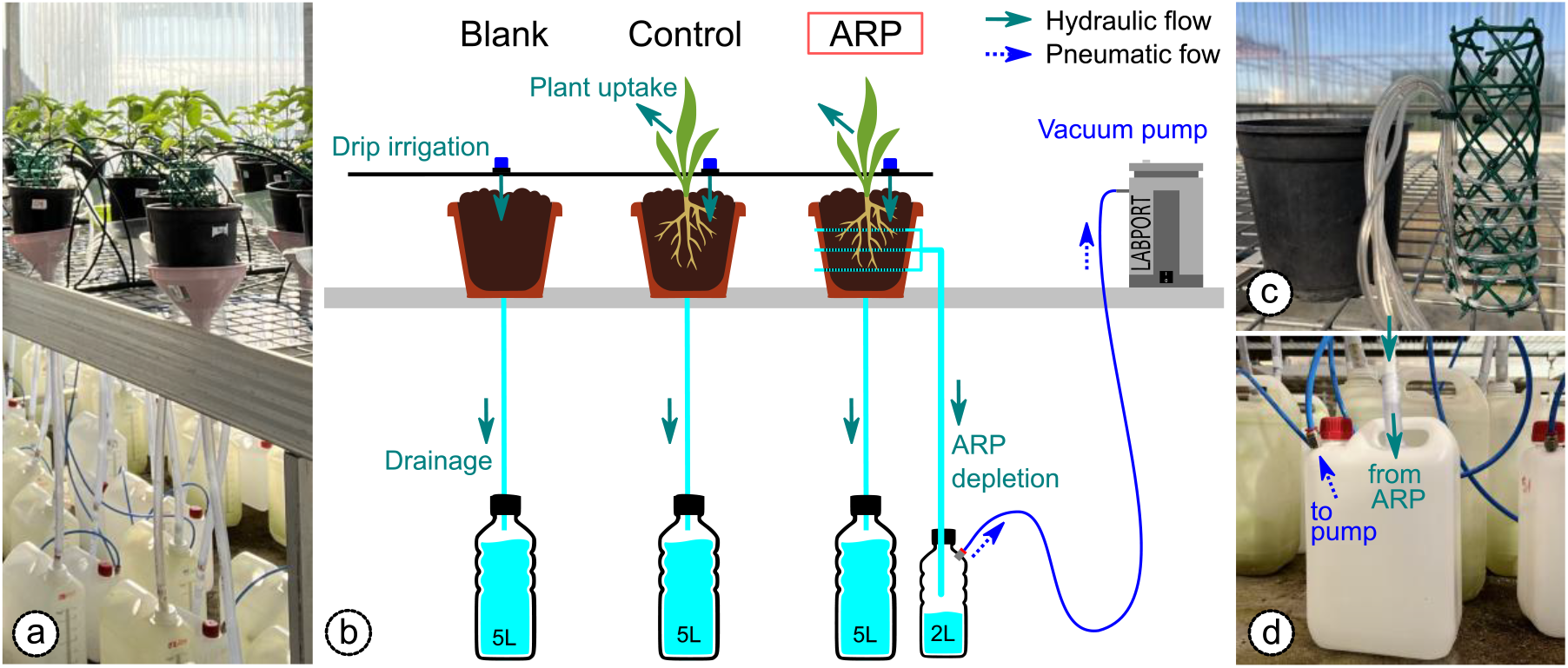
Experimental setup and artificially rooted pot (ARP) system. (a) Photograph and (b) schematic representation of the experimental design. ARPs generate controlled soil water depletion in the absence of living neighbors, thereby isolating exploitative responses from neighbor-derived biological cues. (c) Detail of an artificially rooted pot (ARP). (d) Detail of a 2L vacuum collector container.

## Material and methods

### Model development

We developed a deliberately simple, resource-explicit model to identify the minimal conditions under which soil water depletion can generate contrasting root allocation responses. Although the framework is conceptually applicable to depletable soil resources more broadly, we formulate and test it here specifically for soil water, a dynamic and commonly shared soil resource for which tragedy-of-the-commons logic has previously been developed (Zea-Cabrera *et al*. 2006). Soil water dynamics were represented as a balance between constant input, background abiotic depletion, focal-plant uptake, and neighbor-equivalent depletion by competing roots. Root uptake increased linearly with root biomass and water availability, and root allocation was modeled as a cost–benefit trade-off between water acquisition and the construction and maintenance costs of root biomass. The model is conceptually related to Gersani *et al*. (2001), but incorporates explicit soil water dynamics and isolates depletion from neighbor detection.

Soil water content (w, L) is assumed to vary according to a constant input rate (I, L d^−1^) and losses resulting from abiotic and biotic depletion processes. Abiotic depletion (d_A_, d^−1^) represents losses through processes such as percolation and evaporation, whereas biotic depletion (d_B_, d^−1^) represents water uptake by non-self roots. Soil water uptake by the focal plant depends on root biomass (R, g) and a root uptake coefficient (α, g^−1^ d^−1^). Soil water dynamics are therefore given by

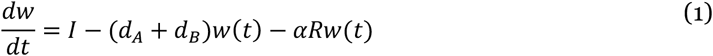

Rather than modeling strategic interactions between competing individuals, we focus on how a single plant adjusts root biomass in response to increasing rates of soil water depletion. Hence, neighboring plants are not modeled explicitly; their effects are incorporated solely through their contribution to soil water depletion. A payoff function (G) for the modeled plant, expressed in water units per unit time, is constructed to determine the optimal root biomass, considering the balance between the benefits of resource acquisition by roots and the costs of constructing and maintaining root biomass (c, L g^−1^ d^−1^). This equation takes the form

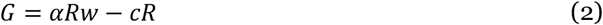

Assuming that water dynamics occur on a much faster timescale than changes in root biomass, water availability can be approximated by its quasi-equilibrium value (dw/dt=0)

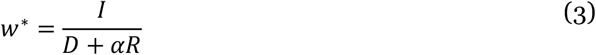

where D = d_A_ + d_B_. Substituting this expression into the payoff function yields

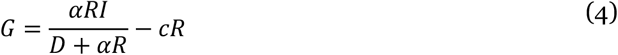

The optimal root biomass R^∗^ is obtained by maximizing G. Since G is strictly concave, as shown by the inequality

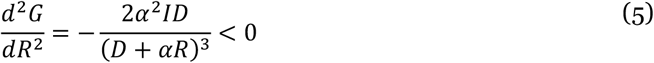

it has a unique stationary point which corresponds to the unique global optimum. Then, the first-order condition for an interior optimum is dG/dR=0. Differentiating G with respect to R gives

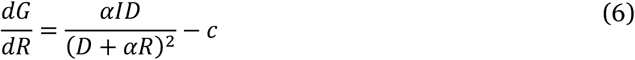

Solving for R yields the optimal root allocation

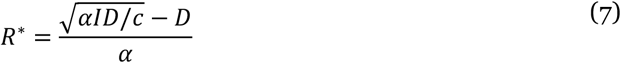

This optimum exists as a positive value only when αI > cD, i.e., the maximum marginal gain from investing in roots exceeds the marginal cost of roots. The optimal root allocation is a concave function of total depletion rate D, but it is not monotonic, as shown by differentiating R^*^ with respect to the total depletion rate D:

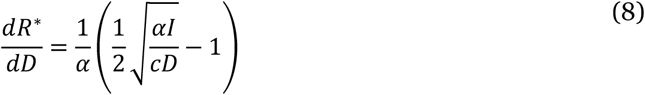

Consequently, increasing soil water depletion does not produce a uniform root response. When depletion rates are low, plants are predicted to increase root allocation as depletion increases. However, beyond the critical depletion rate

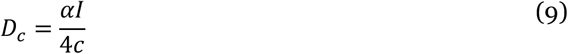

further increases in depletion reduce optimal root allocation. Interpreting D=d_A_+d_B_, this implies that the presence and magnitude of non-self biotic depletion (d_B_) can shift the system across the critical threshold, depending on baseline abiotic depletion (d_A_).

### Greenhouse experiment design

#### Experimental setup

We conducted a controlled pot experiment to quantify plant root allocation responses under varying levels of soil water depletion. Plants were grown either in control 2L pots without artificial depletion structures (CON) or in identical pots equipped with artificially rooted systems (ARPs) designed to impose controlled water depletion rates in the soil volume. In addition, unplanted pots (BLANK) were included to quantify baseline water loss and system performance. In total, 80 experimental units were used: 35 CON plants, 35 ARP plants, and 10 BLANK (**Fig 1**). Seeds of *Capsicum annuum* L. var. *bellotero* were sown on 27 February 2025 in peat-based trays maintained at 30°C. Seedlings were transplanted on 9 May 2025 into pots containing a homogeneous mixture of peat, topsoil, and river sand. Plants were grown for up to 13 weeks, with final harvest on 15 July 2025.

#### Artificially rooted pots (ARPs)

ARPs were constructed to impose controlled soil water depletion via suction-driven extraction. Each system consisted of six perforated PVC microtubing rings (23 cm length, 2 mm internal diameter), connected via T-junctions and distributed throughout the soil volume. Perforations were introduced using a heated needle. Rings were connected to 40 cm non-perforated tubing leading to a sealed 2L collection reservoir. The reservoir was connected via 4-mm pneumatic tubing (Pneufit C fittings) to a Labport N840 variable-flow (20–50 L min^−1^) vacuum pump (KNF Neubergerm, Freiburg-Munzingen Germany) (**Fig 1c, d**). Pump flow rates were adjusted over time: 0 L min^−1^ (weeks 1–2), 30 L min^−1^ (week 3), 40 L min^−1^ (weeks 4–5), and 50 L min^−1^ (weeks 6–13), reproducing the increase in water depletion expected as root systems grow. The system operated daily from 10:00 to 19:00 h simulating the water uptake pattern of C_3_ plants, which absorb water mainly during daylight when stomata are open. Water collected in reservoirs was measured weekly on weeks 9 to 11, and used to estimate effective depletion rates. Observed variation in collected volumes among ARPs was retained and used as a continuous covariate in subsequent analyses.

#### Water supply and drainage

Water was supplied via an automated drip irrigation system operating at 10:00, 13:00, and 16:00 h daily. Pressure-compensated emitters (nominal 2 L h^−1^) were individually calibrated; emitters outside 1.80–2.20 L h^−1^ were replaced. Actual flow rates were used to compute irrigation inputs per pot. Drainage was collected in individual containers beneath each pot via funnels and tubing. Weekly drainage was recorded for all experimental units. Unplanted BLANK pots were used to estimate evaporative losses. Net plant water uptake was calculated in CON pots by subtracting evaporation estimates from planted pots. The saturated water-holding capacity of the substrate in the 2L pots (w_sat_ ≈ 0.73 L) was determined gravimetrically as the difference between the mass of dry substrate-filled pots and the mass of 10 replicate pots after saturation and free drainage. Assuming a water density of 1 kg L^−1^, mass differences were converted to equivalent water volumes.

#### Plant growth conditions and maintenance

Plants were maintained under greenhouse conditions in CULTIVE (URJC, Móstoles). Slow-release fertilizer was initially applied to the pot substrate (Compo Universal NPK with micronutrients, 2.7 g). Because the experiment was designed to focus on plant responses to water dynamics, plants then received weekly foliar fertilization from 21 May onward (Vithal Garden, 10 mL per plant), providing nutrient availability largely independent of soil water dynamics and reducing the likelihood that nutrient limitation drove root allocation responses. Aphid infestations were controlled using pyrethrin in week 2, a commercial insecticide in week 4, and potassium soap washes as needed.

#### Plant measurements and harvest

Plant height and two orthogonal canopy diameters were measured weekly. Plants were harvested between 24 June and 15 July 2025 (weeks 11–13). Shoots and roots were separated, washed, and dried at 60°C for 72 h. Dry biomass was determined using analytical balances.

### Parameterizing the root allocation equilibrium prediction

To obtain measured effective parameters of model Eq. (7), effective parameters I, d_A_, d_B_, and α were estimated from experimental water-balance measurements. Throughout the manuscript, “effective” parameters refer to quantities inferred from the experimental water balance under the assumption that soil water content remained approximately constant. Abiotic losses, ARP-induced depletion, and plant water uptake were quantified independently using orthogonal control and treatment pots, allowing separation of the major components of the water budget. Under quasi-steady-state conditions, the water balance satisfies

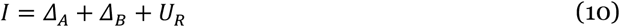

where Δ_A_ and Δ_B_ denote abiotic and ARP-induced depletion fluxes, respectively, and U_R_ denotes root-mediated water uptake. Effective parameter estimates were then used to map experimental conditions onto model predictions of optimal root biomass R^∗^.

#### Abiotic water depletion

Abiotic water loss Δ_A_ (L d^−1^) was estimated from BLANK pots, as the net water balance between water supply (measured from dripping irrigation times and flow) and drainage output:

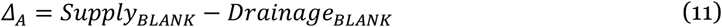

Because unplanted control pots contained no plants or artificial root systems, this flux reflects background water loss due to evaporation and physical drainage processes. Mean abiotic loss was computed across ten unplanted control pots over weeks 1–10 (100 observations total). Δ_A_ was found to have an approximated value of 0.007 L d^−1^.

To obtain the effective abiotic depletion (d_A_, d^−1^), fluxes were normalized by the soil water content (w_sat_), yielding

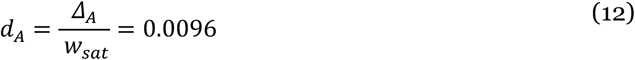

#### Root uptake

Plant water uptake (U_R_, L d^−1^) was estimated for each planted control (CON) pot as the difference between water supply and drainage after accounting for mean abiotic water loss:

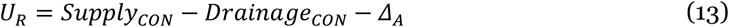

Water supply was calculated from the duration of each irrigation event and the calibrated flow rate of the corresponding emitter.

Root biomass, measured destructively at harvest, was related to canopy volume (V), and calculated assuming a cylindrical canopy geometry from plant height and mean canopy radius, using the allometric power-law relationship

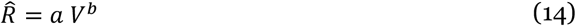

This yields fitted values of a = 0.004043 and b = 0.641169. This relationship was subsequently used to estimate root biomass for every plant at each weekly measurement.

Assuming that soil water content remained approximately constant and close to the saturated water-holding capacity of the substrate (w_sat_), the effective root uptake coefficient (α, g^−1^ d^−1^) was estimated by fitting the mechanistic relationship

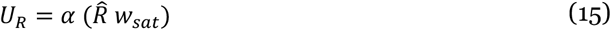

using linear regression constrained to pass through the origin (i.e., with no intercept), consistent with the model assumption that water uptake is zero in the absence of roots. The fitted coefficient was α=0.03199 (adjusted R^2^=0.702, P < 2.2 10^−16^).

#### ARP-induced water depletion

Artificial biotic water loss Δ_B_ (L d^−1^) was assumed equivalent to the volume of water collected in 2L reservoirs connected to the ARP system. Effective biotic depletion rates (d_B_, d^−1^) were obtained by normalizing depletion fluxes by the water-holding capacity of the substrate

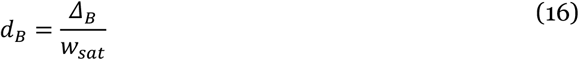

ARP-induced depletion estimates were retained separately for each ARP replicate and used as continuous predictor variables in subsequent analyses.

#### Water input

Because irrigation was applied to saturation, with excess water draining immediately after each irrigation event, irrigation supply did not represent the effective resource input available to plants and was therefore not used directly to estimate the model parameter (I, L d^−1^). Instead, under the assumption that soil water content remained approximately constant through time, the effective resource input was inferred from the daily water balance in equation (10). Under quasi-steady-state conditions, water entering the soil must replace the water lost through abiotic depletion and plant uptake. Accordingly, for control pots

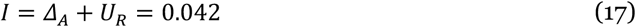

Although effective resource input may have differed slightly among individual pots owing to experimental variability, a single value of I was required for model parameterization. Therefore, the resource input parameter was estimated as the mean value of Δ_A_+U_R_ across all control pots.

## Data analysis

All statistical analyses were performed in R (R Core Team 2026).

### Comparison of root biomass between ARP and control treatments

To determine whether plants grown in Artificial Root Pots (ARP) differed in root biomass from plants grown in control (CON) pots, root dry biomass was compared between treatments. Assumptions of normality were evaluated separately for each treatment using the Shapiro–Wilk test (Shapiro & Wilk 1965), and homogeneity of variances was assessed using Levene’s test (Brown & Forsythe 1974). Root biomass was normally distributed in CON pots (Shapiro–Wilk: W = 0.966, P = 0.340) but not in ARP pots (W = 0.936, P = 0.043), whereas variances did not differ significantly between treatments (Levene’s test: F = 1.348, P = 0.250). Because the normality assumption was violated, differences in root biomass between treatments were assessed using a two-sided Wilcoxon rank-sum (Mann–Whitney) test.

### Empirical analysis of root allocation responses

To evaluate whether root biomass responded to increasing intensity of simulated belowground competition in the direction predicted by the model, root biomass was modeled as a function of the standardized artificial depletion rate, where increasing values of d_B_ represent increasing rates of water depletion by the ARP system. A simple linear regression was first fitted as the null hypothesis of a monotonic response to increasing depletion. A quadratic regression including linear and squared terms of d_B_ was then fitted to represent the unimodal response predicted by the optimization model. Because the linear model is nested within the quadratic model, both models were compared using an analysis of variance (ANOVA).

### Mechanistic model fitting

Having established that the empirical response was consistent with the predicted unimodal pattern, we then fitted the analytical equilibrium solution of the optimization model (Eq. 7) to the observed root biomass using nonlinear least squares. In this formulation, total water depletion is given by D = d_A_ + d_B_, where d_A_ represents background water depletion and d_B_ represents the artificial neighbor-equivalent depletion imposed by the ARPs. The model was fitted by estimating the background depletion rate d_A_, the root water uptake coefficient α, and the input-to-cost ratio K=I/c, such that

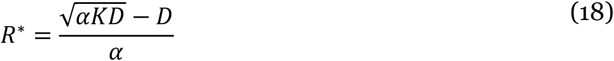

Empirical water-balance measurements were used to obtain biologically informed starting values for nonlinear fitting and to provide benchmarks for interpreting fitted parameters. Because background water loss measured in blank pots may not fully represent the effective depletion environment experienced by planted pots, d_A_ was estimated in the fitted model. The water input rate I was retained as the empirical estimate obtained from water-balance measurements, and the root cost parameter c was subsequently derived from K=I/c.

Mechanistic model performance was compared with the descriptive quadratic model using Akaike’s Information Criterion (AIC) (Akaike 1974). The critical artificial depletion rate at which root allocation reaches its maximum was calculated from Eq. 7 as d_BC_ = αK/4−d_A_. Fitted parameters are reported as estimates ± SE and were compared with empirical water-balance estimates to assess their biological realism (**Table 1**). Because fitted values represent phenomenological parameters describing realized allocation responses, rather than direct physiological measurements of instantaneous water uptake, differences between empirical and fitted estimates were interpreted as informative about the scaling between short-term water-balance processes and final harvested root biomass.

**Table 1:** Water-balance effective estimates and fitted parameters of the mechanistic water-depletion model. Effective values were obtained from empirical water-balance measurements and used as biologically informed starting values or benchmarks for nonlinear fitting. The fitted model estimated the background depletion rate d_A_, root uptake coefficient α, and input-to-cost ratio K=I/c. The root cost parameter c was derived from K using the effective water input rate I shown in the table. The critical neighbor-equivalent depletion rate was calculated as d_BC_=αK/4−d_A_. Fitted values are shown as estimates ± SE; all fitted parameters were significantly greater than zero (P < 0.01).

| Parameter | Meaning | Effective value | Fitted value | Units |
| --- | --- | --- | --- | --- |
| $I$ | Water input | 0.042 | — | $L\ d^{-1}$ |
| $K=I/c$ | Input-to-cost ratio | — | $13.41 \pm 0.77$ | $g$ |
| $c$ | Root cost | — | 0.00313 | $L\ g^{-1}\ d^{-1}$ |
| $d_A$ | Abiotic water depletion | 0.0096 | $0.00296 \pm 0.00111$ | $d^{-1}$ |
| $\alpha$ | Root water uptake | 0.03199 | $0.00244 \pm 0.00049$ | $g^{-1}\ d^{-1}$ |
| $d_{BC}$ | Critical water depletion | — | 0.00522 | $d^{-1}$ |

### ESS-comparable interpretation of the optimization model

To connect our root allocation optimization model to root tragedy-of-the-commons logic, we reinterpreted the imposed neighbor-equivalent depletion rate as the depletion generated by the roots of identical neighboring plants. To express d_B_ as equivalent symmetric neighbor root biomass, we assumed that neighbor roots generate depletion according to the same fitted root-depletion coefficient α:

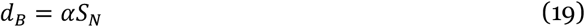

where S_N_ is the total equivalent root biomass of all neighboring plants. Replacing D = d_A_+d_B_ by d_A_+αS_N_ into the fitted allocation function from Eq. 18 gives the focal best-response function B:

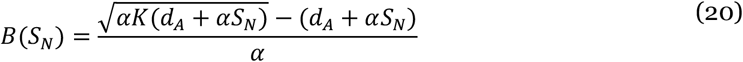

For N identical plants sharing the same resource environment, each focal plant experiences the summed root biomass of its N−1 neighbors. Under symmetry,

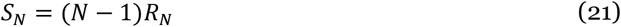

where R_N_ is the root biomass produced by each plant at the symmetric equilibrium. The ESS-comparable equilibrium therefore occurs where

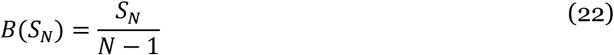

Graphically, this corresponds to the intersection between the best-response curve B(S_N_) and the line from Eq. (22). We solved these intersections numerically using the fitted parameters of the mechanistic model.

We then compared this graphical best-response interpretation with an explicit two-plant version of the water-depletion model, in which the symmetric equilibrium was obtained by solving the first-order conditions for both plants simultaneously (**Supporting Information**). This confirmed that, for two identical plants, the graphical crossing corresponds to the symmetric best-response solution of the explicit two-player model. Because the N=3 and N=10 equilibria are obtained from the same symmetry condition, we used this construction to infer symmetric best-response equilibria for larger numbers of identical competitors.

## Results

Following the conventional approach of comparing plants grown under artificial competition with controls, no significant differences in root biomass were detected between ARP and control treatments (Wilcoxon rank-sum test, W = 595, P = 0.84). However, this categorical comparison ignores the continuous variation in water depletion imposed by individual ARPs. When root biomass was analyzed as a function of the experimentally measured depletion rate, the data revealed a hump-shaped relationship (**Fig. 2**). A linear model detected no relationship between root biomass and depletion (F_1,68_ = 0.43, P = 0.51), whereas a quadratic model provided a significantly better fit (nested ANOVA, F1,67 = 11.18, P = 0.0014) and yielded a significant negative quadratic term (t=−3.34, P=0.0014), consistent with increased root allocation under moderate depletion followed by reduced allocation under stronger depletion.

**Figure 2.**
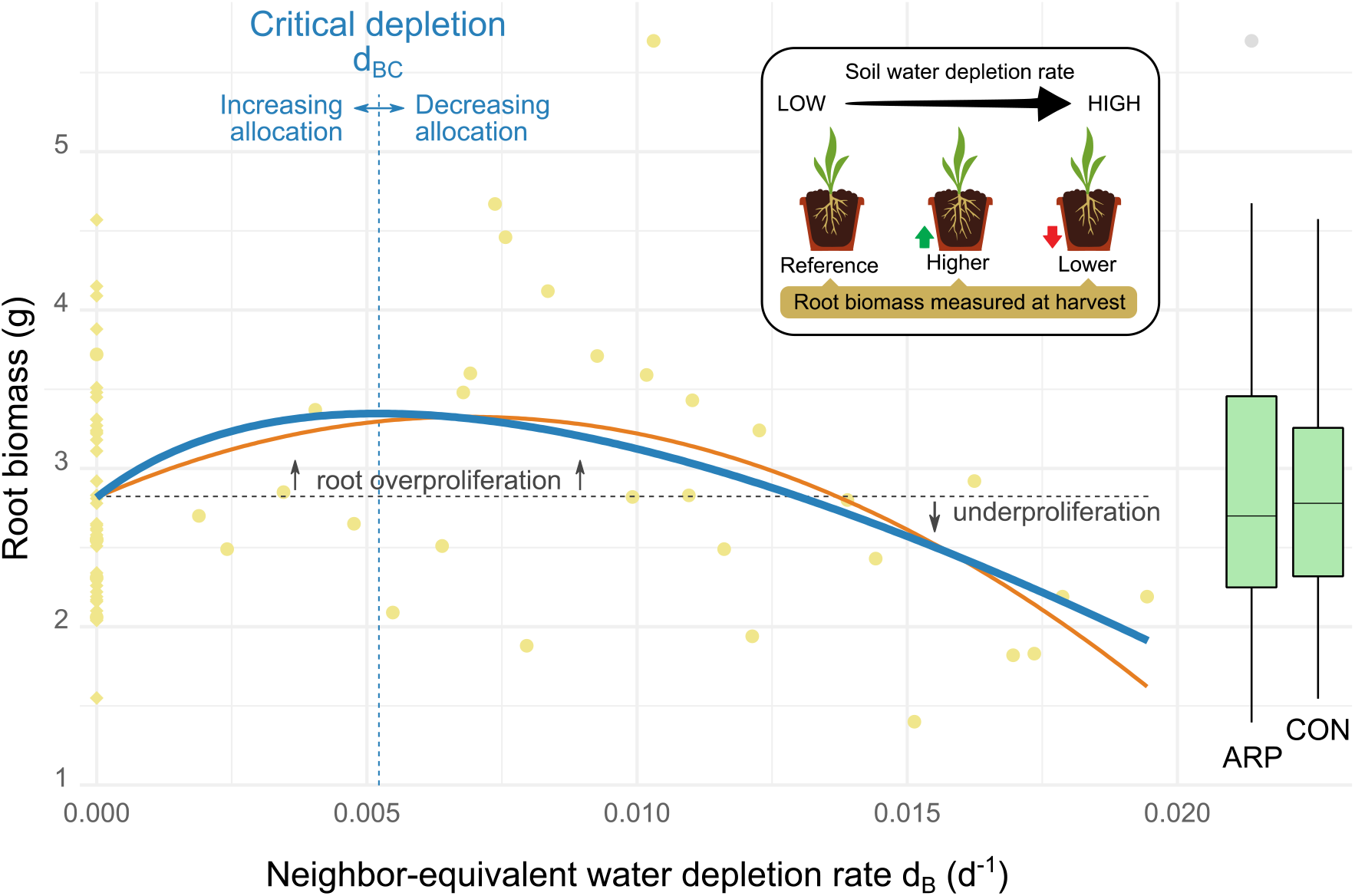
Root allocation responses to increasing neighbor-equivalent water depletion. Root biomass of individual plants as a function of the artificial water depletion rate imposed by artificially rooted pots (d_B_). Points represent individual plants; control plants (diamonds) are shown at d_B_ = 0, whereas ARP plants (circles) span a gradient of depletion rates. The orange curve shows the descriptive quadratic model, and the blue curve the fitted mechanistic model. The blue dashed line indicates the critical depletion rate (d_BC_ = 0.00522) predicted by the mechanistic model, at which the response of optimal root allocation to increasing water depletion changes from positive to negative. The horizontal dashed line indicates the mean root biomass of control plants and is shown as a visual reference for over- and underproliferation relative to controls. Boxplots summarize the distribution of root biomass in the control (CON) and artificially rooted pot (ARP) treatments. Inset schematically illustrates the depletion-dependent root allocation strategy predicted by the mechanistic model and supported by the experimental data.

The theoretical optimization model predicted a non-monotonic (hump-shaped) response of root allocation to soil water depletion, with root biomass increasing at low depletion rates and declining beyond a critical depletion rate. Like the quadratic model, the fitted optimization model (mechanistic) reproduced the observed non-monotonic response, yielding a clear optimum within the experimental range. Its performance closely approached that of the descriptive quadratic model (AIC_mechanistic_ = 164.21 vs. AIC_quadratic_ = 162.88; ΔAIC = 1.34), while preserving the theoretical depletion-response structure. The estimated critical neighbor-equivalent depletion rate was d_BC_ = 0.00522 d^−1^. Fitted parameters were positive and statistically distinguishable from zero (P<0.01; **Table 1**). The fitted uptake coefficient was lower than the empirical water-balance estimate, indicating that fitted parameters should be interpreted as phenomenological parameters describing realized allocation responses rather than direct physiological measurements of instantaneous water uptake.

Using the fitted parameterization, the predicted isolated-plant root biomass was R_1_=2.82 g. The symmetric best-response equilibrium for two plants was R_2_=3.27 g, corresponding to overproliferation relative to plants grown alone, as confirmed by the explicit two-plant ESS solution in the **Supporting Information**. Applying the same symmetric-equilibrium construction to larger numbers of identical competitors predicted R_3_=2.77 g, very close to the isolated-plant prediction, and R_10_=1.10 g. Thus, the model predicts tragedy-like overproliferation relative to isolated plants for two interacting plants, near-neutral responses for three plants, and underproliferation under stronger competitor-driven depletion (**Fig 3**).

**Figure 3.**
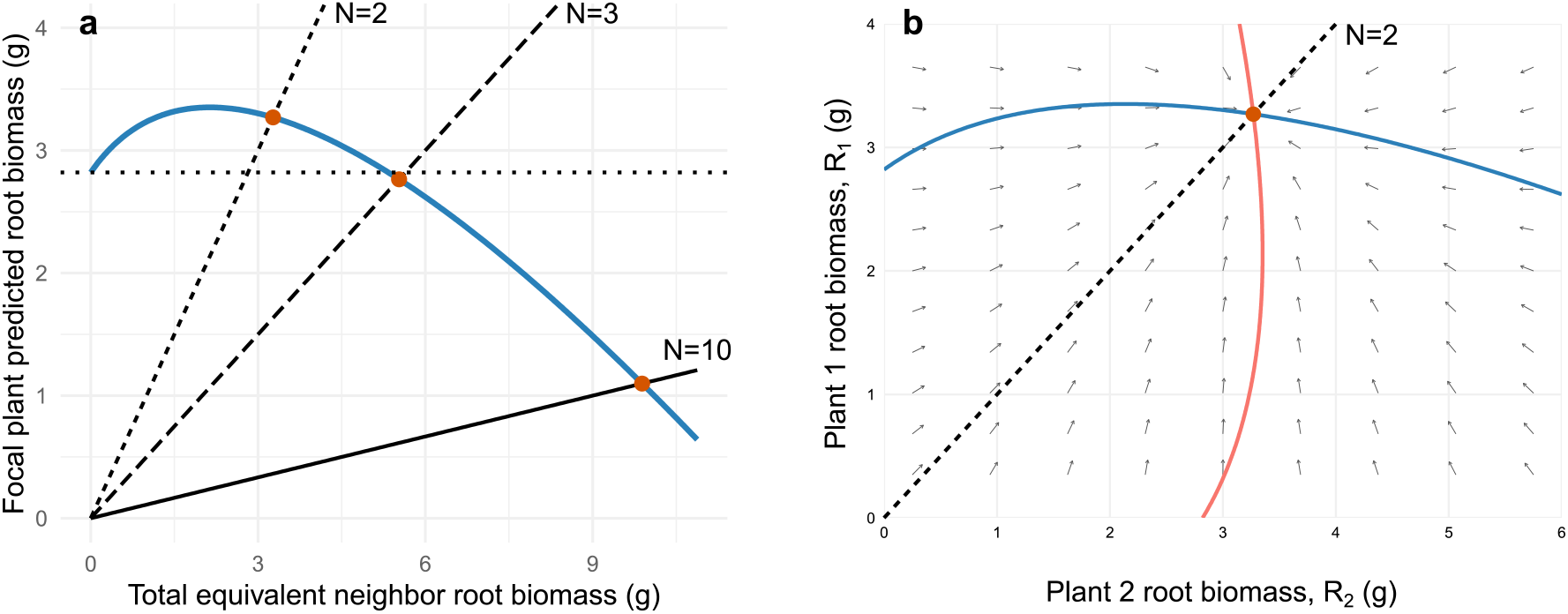
Symmetric best-response interpretation of the depletion-response model. **(a)** Fitted focal best-response function after transforming neighbor-equivalent water depletion into total equivalent neighbor root biomass. Diagonal lines show the symmetric equilibrium conditions for two, three, and ten identical plants sharing a common soil volume. Points indicate the corresponding symmetric best-response equilibria. The horizontal dotted line shows the predicted root biomass of a plant grown without neighbor-equivalent depletion. Under the fitted parameterization, two interacting plants produce more roots per plant than the isolated-plant reference, three plants produce a near-neutral response, and ten plants produce less root biomass per plant. **(b)** Explicit two-plant phase-plane analysis of the same fitted model. Colored curves show the plant-specific isoclines, defined by ∂G_1_/∂R_1_=0 and ∂G_2_/∂R_2_=0. Arrows indicate the direction of simultaneous adaptive change in root biomass expected from each plant’s payoff gradient. The dashed diagonal line indicates equal root biomass in both plants. The intersection of the two isoclines occurs at R_1_=R_2_=3.27 g, confirming the symmetric best-response equilibrium for two identical plants. The analytical derivation is provided in the **Supporting Information**.

## Discussion

By experimentally isolating the depletion component of belowground competition, our study shows that water depletion alone can generate contrasting root allocation responses. This depletion-dependent framework reconciles previously conflicting observations by suggesting that root overproliferation, no detectable response, and underproliferation need not represent distinct strategies, but can instead emerge as different positions along a continuous response to competitor-driven water depletion. Under this framework, plants experiencing moderate rates of water depletion are expected to benefit from allocating more biomass to roots, whereas intense depletion—such as at high plant densities, in the presence of larger neighbors, or in soils where water is rapidly lost through drainage or evaporation—can shift the optimal strategy toward reduced root investment. Thus, the direction of root allocation responses should vary with the rate at which competitors and abiotic soil properties jointly determine the depletion environment experienced by plants. This provides a mechanistic basis for predicting environmental variation in root foraging strategies across both natural plant communities and agricultural systems.

The mechanism underlying this result is simple. Even when water input remains constant, higher depletion rates shorten the residence time of water in the soil, increasing the advantage of rapid capture. Larger root systems can therefore become increasingly advantageous because they intercept a greater fraction of the same water input before it is lost. At the same time, higher depletion rates reduce equilibrium water availability, lowering the returns on further root investment. The balance between these opposing effects generates the predicted non-monotonic allocation response. A critical depletion rate marks the point at which further depletion shifts from increasing to reducing allocation; it is not a universal boundary between overproliferation and underproliferation, which are defined relative to controls.

The symmetric-equilibrium analysis clarifies how our depletion-explicit model relates to the root tragedy of the commons. When neighbor-equivalent depletion is interpreted as the depletion generated by identical neighboring root systems, the explicit two-plant formulation confirms a tragedy-like symmetric ESS: each plant maximizes its payoff given the depletion generated by the other plant, and the resulting equilibrium root biomass exceeds the isolated-plant prediction. However, this result does not extend monotonically to larger numbers of competitors. Extending the same best-response logic to three identical plants produced a per-plant allocation very close to the isolated-plant prediction, suggesting that moderate increases in competitor number may produce little or no detectable root allocation response in empirical studies. For ten identical plants, however, the equilibrium shifted to higher total neighbor-equivalent depletion but lower per-plant root biomass, because intense depletion reduced water availability enough that further root investment became unprofitable. This distinction differs from the original Gersani et al. (2001) framework. In their model, plants competing in a commons share a soil volume that increases with the number of competitors, while space and resources per individual are held constant. Their model predicts increasing root production with competitor number because competition changes the payoff of capturing a share of a common resource pool, and the ESS increasingly weights average rather than marginal returns. In contrast, our model treats competitors as agents of explicit resource depletion. This introduces a second effect absent from the original model: intense depletion can reduce absolute resource availability enough that further root investment becomes unprofitable.

Several limitations define the interpretation and scope of this framework. The fitted parameters are phenomenological descriptors rather than direct physiological measurements, as harvested root biomass integrates uptake, growth, ontogeny, and developmental constraints over the entire experiment, which helps explain the differences between the effective and fitted estimates reported in **Table 1**. The model is intended to test whether the observed allocation response is consistent with its predicted optimization structure, not to provide a physiological parameterization of instantaneous water uptake. Moreover, the framework deliberately isolates exploitative competition and does not imply that neighbor detection, self/non-self recognition, or other biologically mediated responses are unimportant in natural systems (Novoplansky 2019). Likewise, the ARP system and greenhouse conditions simplify the spatial, temporal, and biological complexity of field soils. The model assumptions are most appropriate for rapidly changing resources such as soil water and may also be reasonable for mobile nutrients such as nitrate (Lambers *et al*. 2008), but are less applicable to poorly mobile resources such as phosphate (Hinsinger *et al*. 2011).

## Supporting information

Apendix

## Acknowledgments

This study was supported by the British Ecological Society through a Small Research Grant (TRAgEDy, SR24_1219). C.C. was supported by the Spanish State Research Agency (AEI) through a Juan de la Cierva grant (JDC2022-048613-I). We thank José and Carlos from CULTIVE for their valuable technical advice and support during the experiment.

