## Supplementary material for "Water depletion can generate contrasting root allocation responses to belowground competition": Apendix

### Explicit two-plant ESS equilibrium corresponding to the N=2 best-response result

In the main text, we interpreted neighbor-equivalent depletion as the depletion generated by identical neighboring root systems and used this transformation to compare the fitted depletion-response model with tragedy-of-the-commons logic. For N=2, the graphical intersection between the fitted best-response curve and the 1:1 symmetry line predicted a per-plant root biomass of 3.27 g. Here, we show that this graphical result is not only a visual analogy, but is equivalent to the symmetric solution of an explicit two-plant optimization model. This provides a formal confirmation that the N=2 result satisfies the pairwise best-response condition: neither plant can increase its payoff by unilaterally changing its root biomass while the other plant uses the equilibrium strategy.

For two identical plants sharing the same soil water environment, equilibrium water availability can be written as

$$w^* = \frac{I}{d_A + \alpha R_1 + \alpha R_2} \quad \text{(S1)}$$

where I is water input, dA is background abiotic water depletion, α is the root water uptake coefficient, and R1 and R2 are the root biomasses of plants 1 and 2. The payoff of plant 1 is

$$G_1 = \alpha R_1 w^* - cR_1 = \frac{\alpha R_1 I}{d_A + \alpha R_1 + \alpha R_2} - cR_1 \quad \text{(S2)}$$

and, equivalently, the payoff of plant 2 is

$$G_2 = \alpha R_2 w^* - cR_2 = \frac{\alpha R_2 I}{d_A + \alpha R_1 + \alpha R_2} - cR_2 \quad \text{(S3)}$$

where c is the cost of root biomass production and maintenance.

The optimal root biomass of each plant, given the root biomass of the other plant, is obtained from the first-order conditions

$$\frac{\partial G_1}{\partial R_1} = \frac{\alpha I(d_A + \alpha R_2)}{(d_A + \alpha R_1 + \alpha R_2)^2} - c = 0 \tag{S4}$$

and

$$\frac{\partial G_2}{\partial R_2} = \frac{\alpha I(d_A + \alpha R_1)}{(d_A + \alpha R_1 + \alpha R_2)^2} - c = 0 \tag{S5}$$

At a symmetric equilibrium, both plants produce the same root biomass, so $R_1 = R_2 = R$. The first-order conditions then reduce to

$$\frac{\alpha I(d_A + \alpha R)}{(d_A + 2\alpha R)^2} = c \tag{S6}$$

Using K=I/c, this can be written as

$$\frac{\alpha K(d_A + \alpha R)}{(d_A + 2\alpha R)^2} = 1 \tag{S7}$$

Equivalently,

$$(d_A + 2\alpha R)^2 = \alpha K(d_A + \alpha R) \tag{S8}$$

Solving this equation gives the positive symmetric equilibrium

$$\mathrm{R} = \frac{\alpha K - 4d_A + \sqrt{\alpha K(\alpha K + 8d_A)}}{8\alpha} \tag{S9}$$

Using the fitted parameters of the mechanistic model, α=0.00244, K=13.41, and $d_A$=0.00296, the positive solution is R=3.269288 g.

Thus, the explicit two-plant optimization model gives the same value as the N=2 best-response intersection reported in the main text. This equilibrium corresponds to a maximum of each plant's payoff with respect to its own root biomass. For plant 1, the second derivative is

$$\frac{\partial^2 \mathrm{G}_1}{\partial R_1^2} = -\frac{2\alpha^2 I(d_A + \alpha R_2)}{(d_A + \alpha R_1 + \alpha R_2)^3} < 0 \tag{S10}$$

and the equivalent expression holds for plant 2. Therefore, when plant 2 produces R=3.27 g, plant 1's optimal response is also R=3.27 g, and vice versa. Under the assumptions of this two-plant depletion model, the N=2 graphical crossing is therefore a symmetric best-response equilibrium. Because no plant can improve its payoff by unilaterally changing root biomass while the other plant uses the same strategy, this equilibrium is ESS-comparable in the pairwise sense used here.

The isolated-plant prediction from the same fitted model is

$$R_{alone} = \frac{\sqrt{\alpha K d_A} - d_A}{\alpha} = 2.82 \text{ g} \tag{S11}$$

Therefore, the explicit two-plant equilibrium predicts greater root biomass per plant than the isolated-plant reference:

$$R_{N=2} = 3.27 \text{ g} > R_{alone} = 2.82 \text{ g} \tag{S12}$$

This confirms the main-text interpretation that the fitted depletion-response model produces a tragedy-like outcome for two identical interacting plants: each plant's individually optimal root biomass under competition exceeds the predicted root biomass of a plant grown without neighbor-equivalent depletion. This supplementary derivation does not estimate the collective optimum or extend the analysis to larger N; it simply confirms that the N=2 best-response result in the main text is equivalent to the explicit two-plant symmetric equilibrium.
